# Population-specific imputation of gene expression improves prediction of pharmacogenomic traits for African Americans

**DOI:** 10.1101/115451

**Authors:** Assaf Gottlieb, Roxana Daneshjou, Marianne DeGorter, Stephen B. Montgomery, Russ B. Altman

## Abstract

Genome-wide association studies (GWAS) are useful for discovering genotype-phenotype associations but are limited because they require large cohorts to identify a signal, which can be population-specific. Mapping genetic variation to genes improves power, and allows the effects of both protein coding variation as well as variation in expression to be combined into “gene level” effects.

Previous work has shown that warfarin dose can be predicted using information from genetic variation that affects protein coding regions. Here, we introduce a method that improves the predicted dose by integrating tissue-specific gene expression. In particular, we use drug pathways and expression quantitative trait loci knowledge to impute gene expression—on the assumption that differential expression of key pathway genes may impact dose requirement. We focus on 116 genes from the pharmacokinetic (PK) and pharmacodynamic (PD) pathways of warfarin within training and validation sets comprising both European and African-descent individuals. We build gene-tissue signatures associated with warfarin dose, and identify a signature of eleven gene-tissue pairs that significantly augment the International Warfarin Pharmacogenetics Consortium dosage-prediction algorithm in both populations. Our results demonstrate that imputed expression can improve dose prediction, in a population-specific manner.

## INTRODUCTION

A crucial component to implementing precision medicine is elucidating how genetic variation affects drug response. These gene-drug associations can then be used for tailored drug selection and drug dosing(Ginsburg and McCarthy 2001; Fernald et al. 2011). Genome-wide association studies (GWAS) allow the association of genetic variants like single nucleotide polymorphisms (SNPs) with a drug phenotype. While GWAS have successfully identified thousands of genotype-phenotype associations, there are several limitations (Roukos 2009). Testing a large number of SNPs requires a large study cohort to identify a statistically significant signal. Since SNPs can be population-specific, findings from one population may not be applicable to another population (Rosenberg et al. 2010). Additionally, GWAS often does not identify causal genes (Wang et al. 2010; Ritchie 2012).

Approaches that aggregate SNPs into genes or pathways have been developed to circumvent some of these drawbacks (Limdi et al. 2010; Björkegren et al. 2015). Working within the gene or pathway level typically decreases the number of hypotheses(Wang et al. 2010) and may also bridge population-specific variations measured on different cohorts. Beyond direct measurement of genetic variation, approaches for using measured or imputed gene expression can potentially provide insight into biological mechanism (Li et al. 2013). For example, PrediXcan (Gamazon et al. 2015) imputes the expected baseline expression of a gene based on the allele composition of SNPs in proximity to that gene (cis-SNPs) and uses these predicted expression values to find associations to disease phenotypes.

One mechanism through which SNPs may affect drug response is by modulating the expression level of genes that are key for drug response. These SNP-effects may be population and/or tissue-specific, and the Genotype-Tissue Expression (GTEx) data sets (Consortium 2015) make it possible to assess tissue-specific baseline levels of gene expression for specific ancestries. In this work, we evaluate the degree to which estimation of baseline gene expression can improve estimates of drug response. We use the generic PrediXcan strategy in a modified way: (1) we impute gene expression in a manner that is population-specific; and (2) we impute genes only in tissues where expression quantitative loci (eQTLs) are associated with these genes. In order to have a more interpretable model, we focus on drug pathway genes relevant to the pharmacologic problem (see also (Montgomery and Dermitzakis 2011)). We impute gene expression in specific tissues for each individual using the GTEx compendium (Consortium 2015) (Figure 1) and learn a signature that is predictive of warfarin dose comprised of gene-tissue pairs on a training cohort. We demonstrate the utility of the signatures by predicting warfarin dose in individuals of African American and European descent. Warfarin dose prediction in African Americans is especially challenging, as the currently known genetic variations predict only a small amount of the dose variability (Daneshjou et al. 2013; Drozda et al. 2015). In both populations, our new signatures explain 8%-12% of the unexplained variance in warfarin dose compared to the International Warfarin Pharmacogenetics Consortium (IWPC) algorithm. Our method performs well on African Americans while a generic strategy using PrediXcan does not. Through these improvements, we offer a general approach for prediction of drug response and discovery of associated gene candidates.

**Figure 1.**
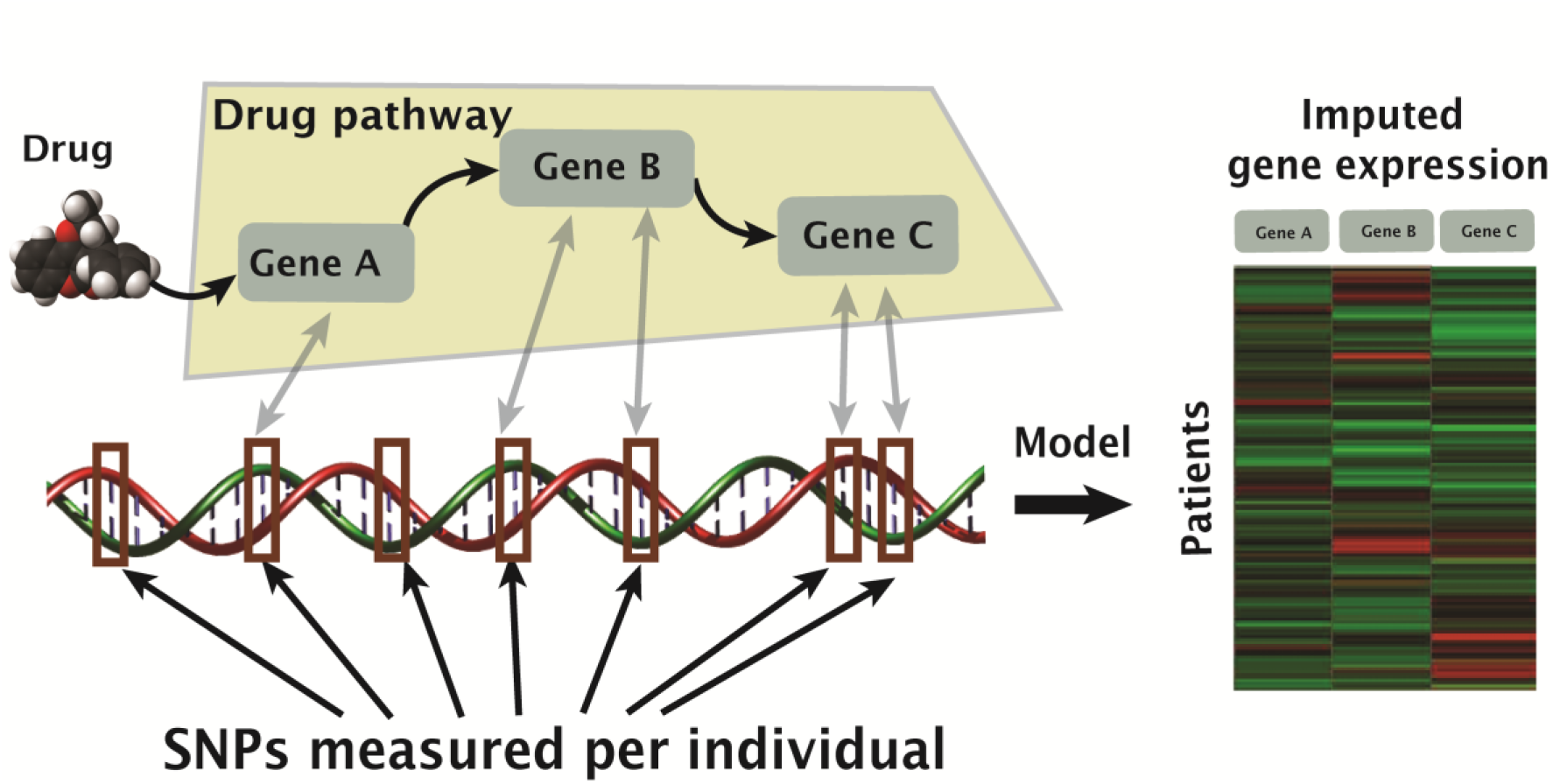
Illustration of the use of SNPs, measured in GWA studies, to impute expression of drug-associated genes.

## RESULTS

Drug response is a complex phenotype and is regulated by multiple genes and across multiple tissues. To identify which genes (in the context of a tissue) might influence warfarin dosing, we (1) imputed the expression of genes in warfarin PD and PK pathways using SNPs in proximity to the gene and (2) learned a tissue-specific gene expression signature. We compared a generic PrediXcan imputation of expression (Gamazon et al. 2015) with a method that (1) builds imputation models that are population-specific and (2) imputes gene expression for each gene only in tissues where the gene has an association with eQTLs (Methods).

We learn the tissue-specific signatures on a European (CEU) and African American (AA) cohorts and validate them by estimating warfarin dose on independent validation cohorts (Methods).

### Imputing tissue specific gene expression

We imputed gene expression by building a regression for each gene-tissue pair using LASSO (Tibshirani 1996). The resulting imputed tissue specific gene expression of PrediXcan (the “generic” strategy) and the “population-specific” method had low correlation across all cohorts (on average, Pearson ρ<0.04, see Methods). Measuring the average correlations between studies, the generic imputations are correlated only between the CEU cohorts while in the population-specific method, both the AA cohorts are correlated and the CEU cohorts (Figure S1, Methods).

### Learning gene-tissue signatures for warfarin dose

We assume that variation in gene expression affects warfarin dose, but this effect might involve only a subset of warfarin pathway genes and could also be tissue-specific. We thus learned a signature, comprised of gene-tissue pairs, that is predictive of warfarin dose using the CEU and AA training cohorts (Methods). We learned a signature with the generic and the population-specific imputation methods separately. The generic CEU signature comprises of sixteen gene-tissue pairs and the population-specific signatures comprise of eleven gene-tissue pairs for the CEU cohort and seventeen gene-tissue pair for the AA cohort (Table 1). Notably, the generic strategy failed to produce a robust signature on the AA training cohort (Methods).

**Table 1.** Predictive signatures for warfarin dose.

| Population-specific |  |  |  | Generic * |  |
| --- | --- | --- | --- | --- | --- |
| African American |  | Central European |  | Central European |  |
| Tissue | Gene | Tissue | Gene | Tissue | Gene |
| Adipose, Subcutaneous | CYP1A1 | Adipose, Subcutaneous | ARRB1 | Adrenal Gland | CCND1 |
| Adrenal Gland | ALOX5 | Adrenal Gland | CUBN | Adrenal Gland | CUBN |
| Brain, Caudate basal ganglia | NCOA1 | Brain, Cerebellar Hemisphere | GGCX | Tibial Artery | JUN |
| Brain, Cortex | ALOX5 | Esophagus, Muscularis | PLCG2 | Brain, Cerebellar Hemisphere | ATF2 |
| Colon Transverse | LGALS2 | Esophagus, Muscularis | UBE2I | Brain, Cerebellum | ATF2 |
| Colon Transverse | STX4 | Liver | VKORC1 | Brain, Cerebellum | GCLM |
| Esophagus, Mucosa | SMAD3 | Skeletal Muscle | PSMA6 | Brain, Frontal Cortex BA9 | AURKA |
| Esophagus, Muscularis | PLCG2 | Skeletal Muscle | PTK2 | Esophagus Gastroesophageal Junction | MGP |
| Heart, Left Ventricle | EPHX1 | Pancreas | LGALS2 | Esophagus Mucosa | CCND1 |
| Liver | VKORC1 | Spleen | GCLC | Heart Atrial Appendage | VKORC1 |
| Tibial Nerve | ITGB1 | Liver | VKORC1 | Liver | CYP2C18 |
| Pancreas | AKT1 |  |  | Pituitary | PSEN1 |
| Pancreas | PLCG2 |  |  | Skin Not-Sun-Exposed Suprapubic | ABL1 |
| Spleen | PROZ |  |  | Testis | SMAD2 |
| Thyroid | EPHX1 |  |  | Thyroid | ITGA2B |
| Thyroid | SERPINF2 |  |  | Thyroid | VKORC1 |
| Whole Blood | GNAI2 |  |  |  |  |

\* The generic methods did not produce a signature on the African American training cohort.

### Predicting warfarin dose

We measured the performance of the signatures based on the R^2^ of the unexplained portion of the IWPC algorithm and by how significant it is relative to shuffled and to random signatures’ backgrounds, correcting for false discovery rate across signatures (Methods).

#### Signatures for patients of European descent (CEU)

Both the generic and the population-specific strategies produced signatures that performed better than the background, with the generic strategy obtaining higher R^2^ results on the CEU validation set (R^2^=0.2, p<0.008 and R^2^ =0.08, p<0.05 for generic and population-specific strategies, respectively, Table 2, Figure 2A). The population-specific signature displayed good performance on the AA validation set (R^2^ = 0.09, p<0.02, Table 2, Figure 2A).

**Table 2.** Performance of the generic and population-specific signatures on different warfarin studies. P-values below false discovery rate of 0.05 are bolded.

| Validation Cohort | Signature | $R^2$ Regression against IWPC residuals | P-value of difference from the background * |
| --- | --- | --- | --- |
| CEU validation | CEU, Generic | 0.2 | **0.008** ($e^{-4}$) |
|  | CEU, Population-specific | 0.08 | **0.004** (0.05) |
|  | AA, Population-specific | 0.1 | **0.004** (**0.03**) |
| AA validation | CEU, Population-specific | 0.09 | **0.0009** (**0.02**) |

\* Background computed as random signatures. P-values of shuffled signatures in parentheses.

**Figure 2.**
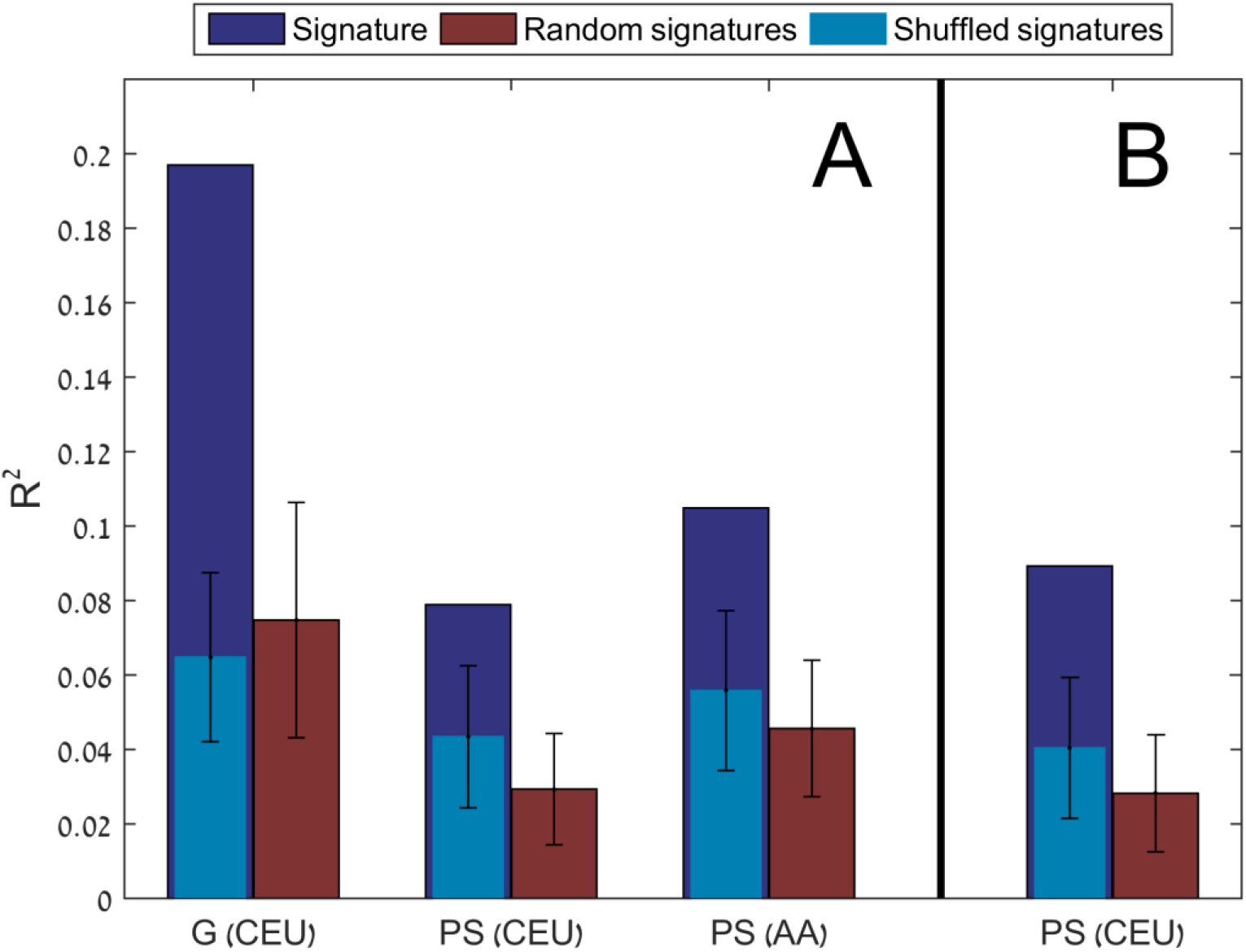
R^2^ results of the predicted unexplained variance in warfarin dose by the IWPC algorithm for the CEU (A) and AA (B) validation cohorts. Represented are the signatures (dark blue), random signatures (purple) and signatures on shuffled data (light blue) as the background models for the AA and CEU signatures. CEU and AA in parenthesis are the training cohort for the signature, G = generic imputation method and PS=population-specific imputation.

#### Signatures for patients of African American descent (AA)

The generic imputation strategy failed to produce a robust signature on the AA training cohort and the population-specific AA-trained signature was not significantly better than the background on the AA validation set. However, the population-specific AA-trained signature was significant on both the CEU validation and CEU training cohorts (R^2^=0.1, p<0.01 and R^2^ =0.24, p<0.0001 for the CEU validation and training, respectively, Table 2, Figure 2B).

### Analysis of gene-tissue pair signatures

Three tissue-specific gene expression levels are correlated with warfarin dose in the CEU validation cohort, but none significant in the AA validation cohort: (1) VKORC1 in liver and in thyroid (Pearson correlation, ρ=0.49, p<e^−15^ in liver and ρ=0.28, p<e^−5^ in thyroid); (2) STX4 in transverse colon (ρ=0.3, p<2e^−6^); and (3) CYP2C18 in liver (ρ=0.35, p<3e^−8^). Out of these three, VKORC1 in liver is the only single predictor of dose that is better than the background (R^2^ = 0.03, p<0.005 for the population-specific imputed expression and R^2^ = 0.03, p<0.02 for the generic).

Besides VKORC1, two genes are common to the CEU and AA population-specific signatures (Table 1): PLCG2 in muscularis esophagus and LGALS2 in pancreas (CEU signature) and transverse colon in (AA signature). CUBN in adrenal gland is common to the CEU generic and CEU population-specific signatures.

## DISCUSSION AND CONCLUSIONS

We have introduced a method to impute gene expression in the context of drug pathways and to select a signature of gene-tissue pairs that are associated with drug response. By focusing on drug response-associated genes, we increase the likelihood of finding biologically relevant variants that influence transcriptional regulation in the context of a drug. We compared the current state of the art, PrediXcan, to a modified, population-specific method on cohorts of patients on warfarin of African-Americans or European descent.

The generic strategy for imputation better explained warfarin dose than our modified method on individuals of European descent, but performed poorly on African Americans. This is not surprising, given the preponderance of European-descent individuals used to generate the imputation model. Indeed, the population sampled in GTEx includes 84% white and only 14% African Americans. Thus, imputing gene expression without the context of the cohort is likely biased towards European populations. The population-specific method, on the other hand, generalized well also to African Americans, with European-trained signatures performing well on African Americans and vice versa. The results are thus consistent with previous findings that warfarin dose models trained on individuals of European descent have poor performance on African Americans (Roper et al. 2010; Scott et al. 2010; Shaw et al. 2010; Daneshjou et al. 2013). Our results suggest that population-specific models are advantageous in cases where the set of SNPs measured for the cohort differ from the set used for building the models and calls for diversifying the sampled populations in GTEx in order to produce better pharmacogenomic models. As the gender and age distributions in GTEx (34% females and majority of individuals 50-70 years old) differ from the distributions in the warfarin cohorts (Table 3 and Figure S2), we estimate that models that further stratify the GTEx population based on these covariates may produce more accurate imputation models and improve the pharmacogenomics models.

**Table 3.**
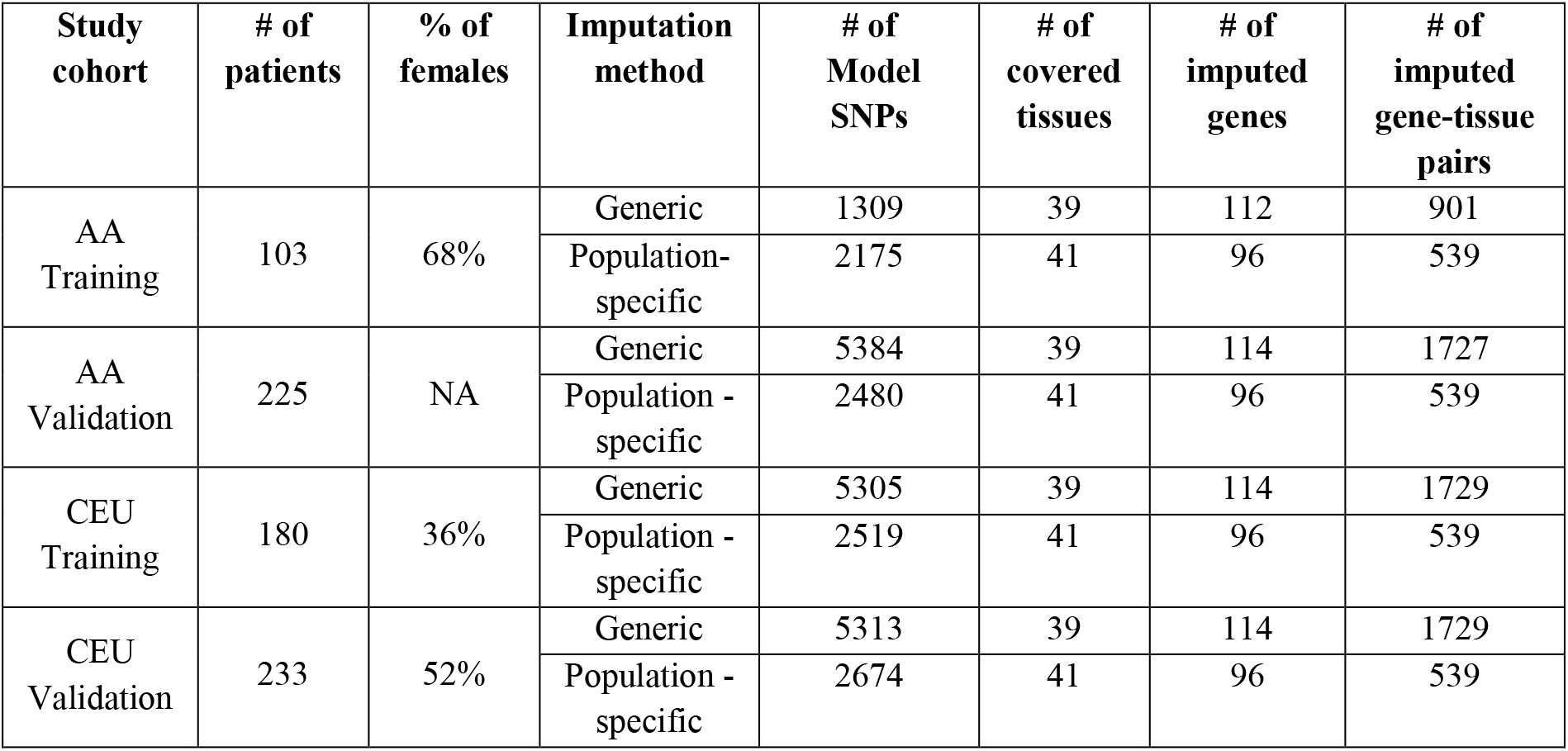
Cohort statistics.

Warfarin acts as an inhibitor of VKORC1 and is part of the IWPC dose algorithm. Polymorphisms in its cis-SNP rs9923231 account for approximately 25% of the variance in stabilized warfarin dose and is currently considered the single largest predictor of warfarin dose (Owen et al. 2010). Notably, the models for both the AA and CEU training cohorts does not include this SNP, nor do they include rs61162043, a SNP that was associated with African American population (Hernandez et al. 2014) (including 13-19 different SNPs in generic model and 3-13 different cis-SNPs in the population-specific models, depending on the cohort). It is thus encouraging that imputed expression of VKORC1 in liver was still found to be a strong dose predictor, suggesting that a significant component in VKORC1 effect on dose is through transcriptional regulation.

Only three genes, VKORC1, STX4 and CYP2C18, are individually correlated with warfarin dose, accounting together for less than a third of the entire signature explained dose (R^2^=0.035, p<0.02 on the CEU validation cohort and insignificant on the African American cohort), which suggests that gene expression associated with dose is combinatorial in nature and supports our pathway-directed multivariate analysis methodology. Additionally, the population-specific signatures include genes with known polymorphism associated with warfarin dose such as GGCX, within African Americans (Cavallari et al. 2012) and EPHX1 within Caucasians (Liu et al. 2015). Lastly, CYP2C18, appearing in the generic signature, was previously reported to be associated with warfarin dose (Wadelius et al. 2007). The report explained this association by linkage disequilibrium of CYP2C18 SNP rs7896133 with CYP2C9*3 (CYP2C9 has no model in liver in PredictDB). rs7896133, is indeed moderately linked to CYP2C9*3 (r^2^=0.68, using SNAP tool (Johnson et al. 2008)) and is included in PredictDB models for CYP2C18 in liver but the generic models include additional eight cis-SNPs, four of them (rs7067881, rs9332214, rs7920801 and rs7088784) with similar or larger weights than rs7896133 and only two of these in LD with CYP2C9*3 (rs9332214 and rs7088784, r^2^=0.87 and 0.68, respectively), suggesting that CYP2C18 might have another mechanism of association with warfarin dose.

Liver is associated with two (out of three) gene-tissue pairs correlated with warfarin dose (VKORC1 and CYP2C18). Liver indeed plays a role in the metabolism of warfarin (Rost et al. 2004). We have included all tissues and have not weighted them based on relevance to warfarin response. Thus, some of the tissue expression we use are from tissues not typically considered relevant to warfarin’s mode of action. These expression values may correlate with expression in other tissues or may represent evidence of an unexpected role of new tissues in warfarin response. Tissue-specific knowledge may improve our methodology and should be considered in follow-up work, taking into account also tissue-specific detection sensitivity (Ardlie et al. 2015).

In conclusion, imputation of the expression of genes relevant to drug action can increase the power of GWA studies to explain drug response. Focusing on genes with genetically driven (versus environmentally driven) expression allows us to build models of drug response that reflect the expected expression levels of critical genes in particular tissues. We have further shown that these models can be population-specific to improve predictive power.

## METHODS

### Datasets

#### Expression Data

Gene expression and eQTLs associated with 42 tissues were extracted from GTEx consortium version 6 (Ardlie et al. 2015) (excluding cell-lines of EBV-transformed lymphocytes and transformed fibroblasts. Tissue statistics are available on the GTEx portal, http://www.gtexportal.org/home/tissueSummaryPage).

We imputed the expression of 116 genes that comprise the curated warfarin pharmacodynamic (PD) and pharmacokinetic (PK) pathways from PharmGKB (Whirl-Carrillo et al. 2012), and the predicted warfarin PD pathway genes (Gottlieb and Altman 2014).

#### Training and Validation Cohorts

We selected a signature, comprised of gene-tissue pairs predictive of warfarin dose, using a training cohort and validated the signature performance in predicting warfarin dose on a validation cohort (Table 3 lists cohort statistics and Figure S2 cohort age distribution). The training set for the African American (AA) signature comprised of 103 previously exome-sequenced individuals of African American descent which received either low dose (<=35 mg/week) or high dose (>49 mg/week) warfarin (Daneshjou et al. 2014). The Central European (CEU) training cohort is the Cooper *et al.* dataset with 180 genotyped individuals (five of which of Hispanic origin) (Cooper et al. 2008). Validation was conducted on 225 genotyped individuals of African American descent (Daneshjou et al. 2013; Perera et al. 2013) and 233 genotyped individuals of European descent (Isma et al. 2009) (see Figure S3 for distribution of doses in each study). The warfarin patient cohorts included 0.02%−1.1% of missing values, which were imputed using *k*-nearest neighbors impute (*k*=5). Using 77 African American individuals mutual to the training exome-sequenced cohort and the genotyped validation cohort, we estimated the measurement error rate as 1% allelic mismatches (including missing values). These 77 individuals were subsequently excluded from the validation set.

### Imputing tissue specific gene expression

In accordance with PrediXcan methodology(Gamazon et al. 2015), imputing gene expression was restricted to *cis*-SNPs, defined as closer than 1 Mb to the outer bounds of the gene (gene and SNP positions extracted from the human genome reference sequence GRCh38 and dbSNP build 144). We focused on gene-tissue pairs where the gene had an associated eQTL (q-values <=5%) in GTEx (Consortium 2015) resulting in 67,022 SNPs in cis with the warfarin-pathway genes that were also measured in at least one of the four warfarin studies used for training and validation.

For the generic PrediXcan imputation, we used the software package of PrediXcan with the weighted *cis*-SNPs in the PredictDB database (Wheeler et al. 2016) to generate imputed gene-tissue pairs. For the population-specific strategy, we imputed each gene only in tissues where the gene has at least one significant eQTL. As the warfarin GWA studies of central European (CEU) population (Cooper et al. 2008) and African American (AA) populations (Perera et al. 2013) measured only a partial set of the SNPs available in GTEx, our population-specific strategy was to build a model per cohort, based only on the SNPs measured in that cohort study (Figure 3A). The gene expression regression models where computed with LASSO (Tibshirani 1996) using five-fold cross validation to select the optimal regularization parameters. Between 111 and 114 genes were imputed using the generic PrediXcan and 96 genes were imputed using the population-specific method (Table 3). Figure S4 display the relative overlap of *cis-* SNPS measured in each warfarin study that were used in imputing the genes.

**Figure 3.**
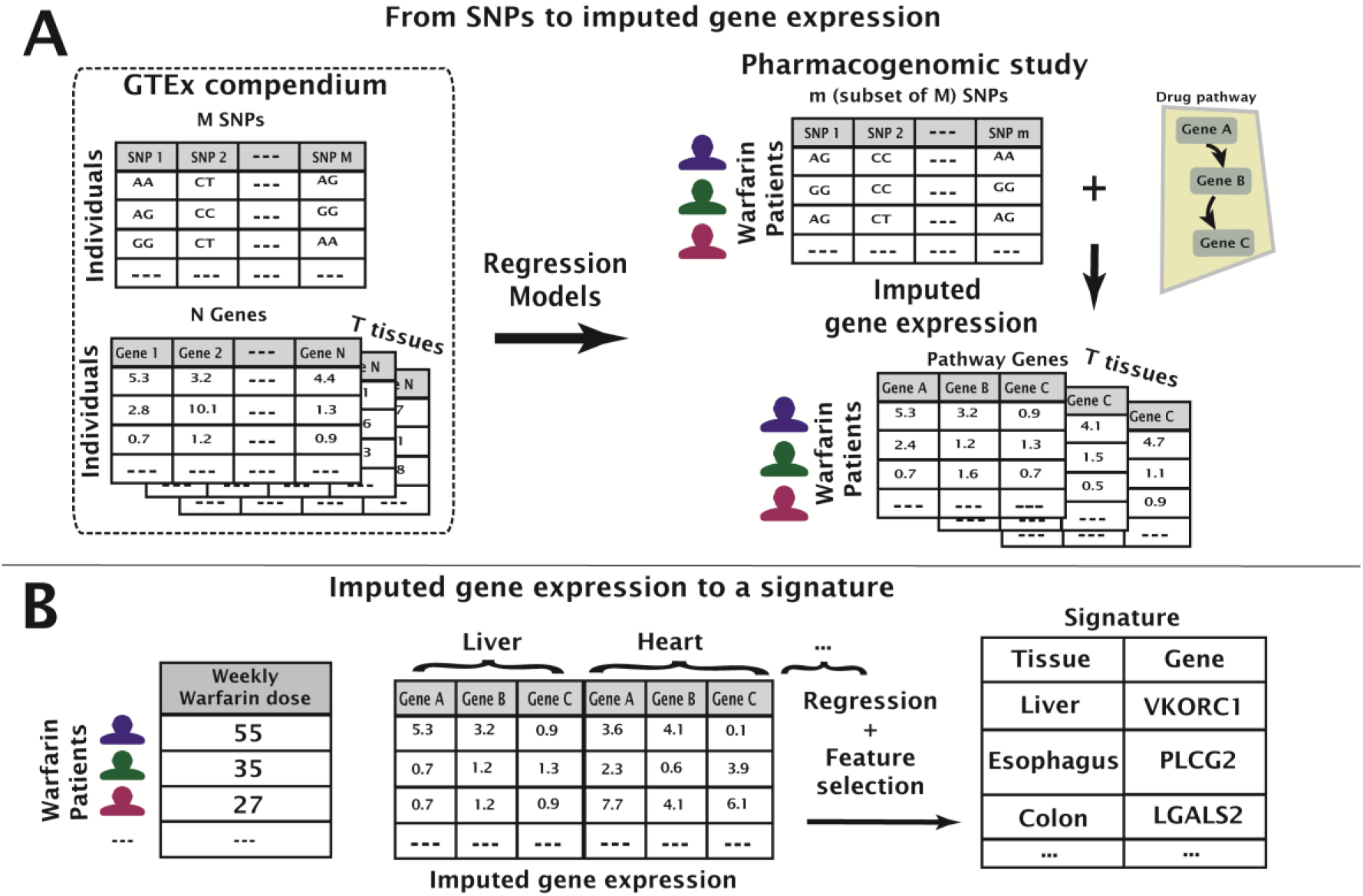
Illustration of the feature construction and signature selection methods. First, gene expression is imputed by regression models from cis-SNPs (A). Then, a signature is learned by regressing the drug response on the imputed expression features (B).

We evaluated the imputed gene-tissue expression in two ways: (1) overlap of imputed genes across the four warfarin study cohorts for each imputation method independently (between-study similarity); and (2) the consistency of each study cohort across imputation methods (between-method similarity). For the between-study similarity, we computed the mean value of each gene-tissue pair across the individuals in each study cohort and computed the Pearson correlation between those means across all gene-tissue pair across the cohorts. For the between-method similarity, we computed the correlations of each gene-tissue pair across individual patients and averaged across all gene-tissue pair. All computations were performed on MATLAB v8.4.

### Learning signatures for warfarin dose

Gene-tissue signatures were learned using LASSO regression analysis with five-fold cross validation, selecting the shrinkage parameter which provided the minimal mean square error (Figure 3B). We repeated this procedure a hundred times with different random cross-validation partitions and selected for the signature gene-tissue pairs appearing in more than half of the repeats.

### Predicting warfarin dose

To validate the signatures’ utility, we computed the predicted dose according to the dosing algorithm published by the International Warfarin Pharmacogenetic Consortium (IWPC) for each validation cohort and inferred the “unexplained dose” defined as the difference between an individual’s actual therapeutic dose and the IWPC predicted dose. In the AA validation dataset, the IWPC dose was computed without the information regarding use of enzyme reducers (phenytoin, carbamazepine or rifampin), which could potentially change the predicted dose by up to 9% (Perera et al. 2013). We performed linear regression using the signatures against the “unexplained dose”. Each signature’s performance was compared to two background models: (1) a shuffled model, in which the “unexplained dose” was shuffled 10,000 times; and (2) a random signatures model, created from 10,000 equal-sized randomly selected signatures, chosen from the gene-tissue pairs not in the signature. P-values of regression R^2^ values were empirically computed relative to these background models and corrected for false discovery rate (FDR) (Benjamini and Hochberg 1995) of 0.05 across the tested signatures.

## ACKNOWLEDGMENTS

### Funding

R.B.A and A.G are funded by the NIH grants LM05652, GM102365 and GM061374. RD is funded by Stanford Medical Scientist Training Program and the Paul and Daisy Soros Fellowship for New Americans. MKD is funded by a Banting Postdoctoral Fellowship. SBM was funded by the Edward Mallinckrodt Jr Foundation and NIH grants R01MH101814, R01HG008150, U01HG007436 and U01HG00908001.

## AUTHOR CONTRIBUTIONS

AG conceived the paper, performed the experiments and analyzed the data. RD and RBA contributed to the experimental design. MKD and SBM computed the eQTL associations on the GTEx data. AG, RD, MKD, SBM and RBA wrote the manuscript.

## Supplementary Figures legends

**Figure S1**: Relative overlap of measured cis-SNPs between cohorts (Jaccard score).

**Figure S2**: Age distribution for each of the warfarin study cohorts.

**Figure S3**: Weekly dose distribution or each of the warfarin study cohorts.

**Figure S4**: Pearson correlations between imputed gene expression means across cohorts for the generic and population-specific imputation methods.

